# Carbon nanodots as a red emissive fluorescent probe for the super-resolution microscopy of DNA dynamics during paclitaxel treatment

**DOI:** 10.1101/2024.10.29.620846

**Authors:** Richa Garg, Kush Kaushik, Abdul Salam, Runmi Kundu, Shilpa Chandra, Chayan Kanti Nandi

## Abstract

Paclitaxel is a commonly used frontline chemotherapeutic drug for cancer treatment. It is known to be functional by arresting the microtubule disassembly during mitosis. Recently, a non-mitotic pathway has been evolving and thus contemplating the mitotic mechanism. Here, using super-resolution microscopy (SRM), we directly visualized the nuclear dynamics and unveiled the mechanism of paclitaxel treatment. A new class of non-toxic, biocompatible and highly fluorescent carbon nanodots (CNDs) were used as a fluorescent probe that were highly capable to directly stain the nuclear DNA, capture the SRM imaging of chromosome and the chromatin structures. Apart from SRM imaging of chromosomes during all stages of normal mitotic cell division, CNDs successfully visualized the formation of lagging, mis-segregated and bridging chromosomes which leads to the multi-micronucleus formation upon paclitaxel treatment. A detailed chromatin remodelling analysis suggested that heterochromatin played an important role in the formation of condensed multi-micronucleus, ultimately leading to cell death.

**Graphical Abstract:** 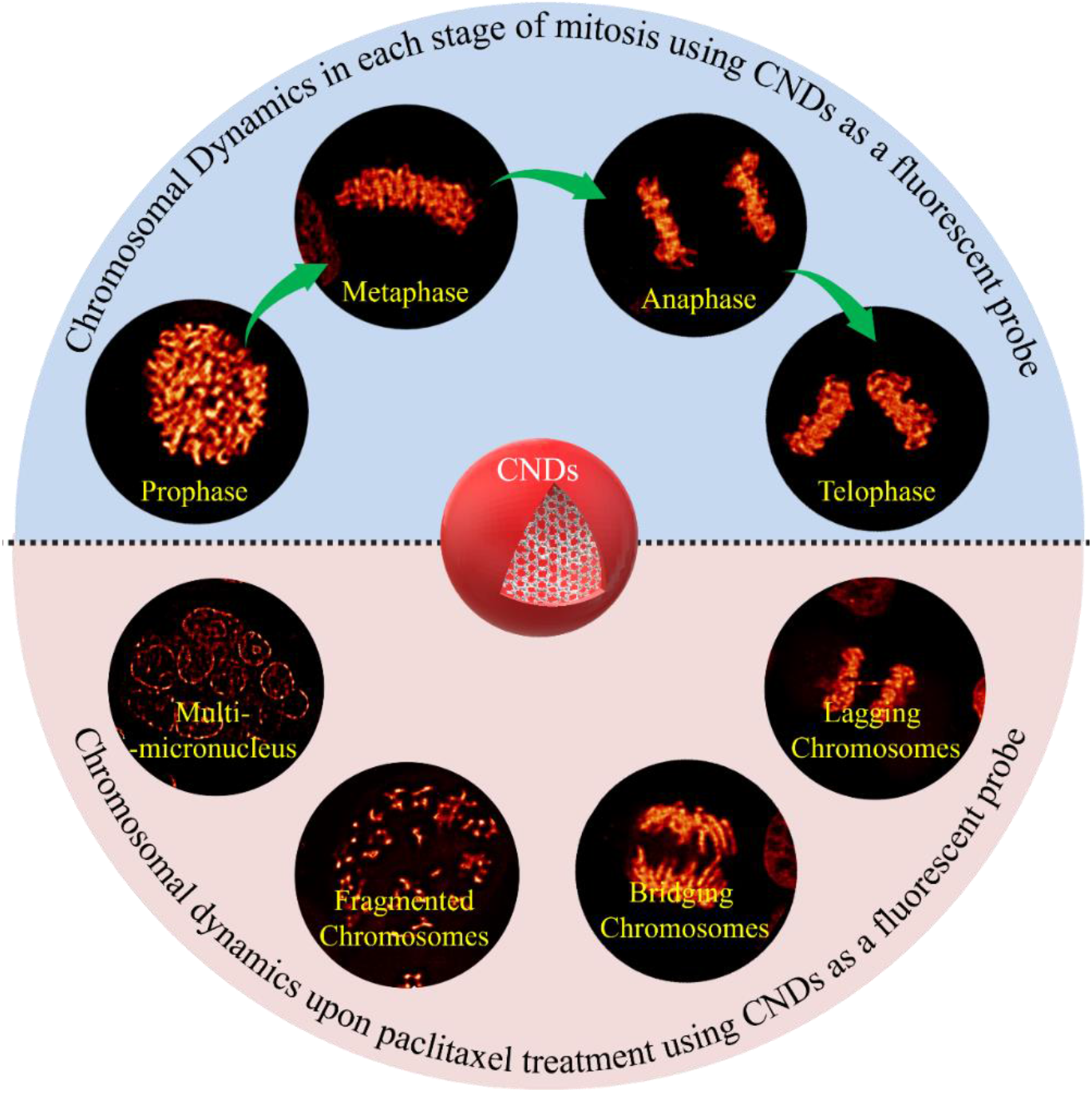

## Introduction

Paclitaxel, a frontline antimitotic chemotherapeutic drug belongs to the taxane family has been extensively used in cancer treatment^[1]^. Despite the impressive clinical success, still a major challenge remains is the development of drug resistance in recurrent cancer as well as reducing the side effects of paclitaxel including heart problems, increased lung inflammation, birth defects, shortness of breath and several other related effects^[2,3]^. These problems may be overcome by knowing the exact mechanism of its treatment and action pathways. A generally accepted mechanism of paclitaxel action is the binding and stabilizing of the beta subunits of microtubules during mitosis and arrest of the mitotic pathway^[4,5]^. Such a proposed mechanism has also been found to be dependent on the therapeutic concentration of the drug, which leads to the alteration of the side effects as well^[6–9]^. As a result, it is essential to accurately understand the mechanism of paclitaxel treatment to overcome such problems.

Recently, a non-mitotic pathway of paclitaxel treatment is evolving, where paclitaxel-bound microtubules exert physical force and distort the nuclear envelope by linking the nucleus and cytoplasm bridge^[1]^. This connects microtubules as well as nuclear lamina. As a result, the formation of the multi-micronucleus occurs in mammalian cells when chromosomes lag during anaphase or fail to align on the metaphase plate^[10]^. Chromosomes that fail to segregate during the interphase of cell cycle, result in a separate nuclear envelope around mis-segregated chromosomes, thus forming their own nuclear envelope known as multi-micronucleus^[11]^. Studies have shown that multi-micronucleus have reduced functioning as compared to the primary nucleus and micronucleated chromatin often undergoes extensive damage in interphase^[12]^. As a result, understanding the exact mechanism of paclitaxel treatment, still remain an open question that needs serious consideration for the successful future of the drug by reducing its side effects ^[13–16]^.

We have utilized a unique approach to understand the detailed mechanism of paclitaxel by using SRM, which is a powerful technique to directly visualize the cellular organelle and their dynamics down to the nanometer level ^[17–20]^. The SRM technique has recently been revolutionized by breaking the diffraction limit of light and provide very insightful ultrastructural dynamical information of various unknown biological events ^[21]^. The SRM technique often requires a suitable fluorescent probe with the desired optical properties such, as high quantum yield (QY) and brightness, along with biocompatible and reduced cytotoxicity^[22]^. Many natural and synthetic fluorescent probes, such as organic dyes^[23]^, fluorescent proteins^[24]^, and quantum dots^[25]^ are available for SRM imaging. However, the organic dyes often suffer with photostability and rapid photobleaching^[26]^. On the other hand, despite the desired optical properties for SRM imaging, quantum dots suffer from the toxic effect on the cellular environment^[27]^. Fluorescent protein with their large size, tedious to obtain and low brightness, limited their applications in SRM imaging. Thanks to the recent development of a new class of highly fluorescent CNDs which exhibit their excellent biocompatibility, very low toxicity and easy synthesis make them an efficient probe for bioimaging of various cellular organelles without disturbing their structure^[28–32]^. The smaller size of ∼ 3-5 nm with their tunable emission characteristics even allows them to be a suitable candidate for SRM imaging^[33]^.

Here, by synthesizing red emissive water-soluble CNDs with direct staining capability to the nuclear DNA, we provide very insightful information on chromosomes and chromatin dynamics in each step during the normal mitosis process and upon paclitaxel treatment. The formation of lagging, bridging and mis-segregated chromosomes that leads to the multi-micronucleus formation during paclitaxel treatment were visualized using super-resolution radial fluctuation (SRRF) microscopy. The results revealed proper chromosomal structure under normal conditions inside the nucleus, but paclitaxel-treated nuclei exhibited unique deformed morphology due to the deformed inner nuclear membrane along with fragmented chromosomes, finally leading to cell death. A detailed chromatin remodelling analysis suggested that heterochromatin played a vital role during paclitaxel treatment.

## Result and Discussion

CNDs were synthesized via a hydrothermal method using glutathione (GSH) and o-phenylenediamine (OPDA) as precursor molecules. In brief, 100 mg OPDA and 90 mg GSH were dissolved in 10 ml of deionized water (DI) and 5% hydrochloric acid (HCl) solution. Subsequently, the reaction mixture was sonicated for 15 minutes and placed into a 25 ml hydrothermal reactor. After hydrothermal treatment, the resultant solution mixture was allowed to cool at room temperature. Further, the resultant reaction product was centrifuged at 10000 rpm for 20 minutes to remove undesired impurities from the reaction mixture. Dialysis was then performed against DI water to obtain a pure sample of CNDs. The solid sample was then obtained by freeze-drying. The obtained CNDs were systematically characterized and then used for further biological study. The as-synthesized CNDs appeared greenish blue when observed to the naked eye and they showed red color upon 488 nm laser excitation.

The transmission electron microscope (TEM) analysis suggested the average size of the CNDs is 4.6 nm (**Figure 1a**). The Raman spectroscopy **(Figure 1b)** showed two broad peaks at 1370 cm^-1^ and at 1557 cm^-1^, which are typical signatures of the D band originating from disordered or glassy carbon and the G band of graphitic in-plane vibrations of sp^2^ bonded carbon atoms in a two-dimensional hexagonal lattice^[34]^. The powder X-ray Diffraction (P-XRD) measurement showed a broad peak at around 24° **(Figure 1c)**, which is due to 002 planes of defective graphitic nature and is attributed to the amorphous nature of CNDs ^[35]^. A few sharp peaks were visible on the top of the broad peak suggesting some crystalline nature of the CNDs. The surface properties of the CNDs were understood by the Fourier transform infrared spectroscopy (FTIR) and X-ray photoelectron spectroscopy (XPS) measurements. The broad band from 3719 cm^-1^ to 2134 cm^-1^ in FTIR was attributed to the stretching vibrations of -OH, NH_2_, =C-H and S-H groups (**Figure S1**). The peak at 1755 cm^-1^ is attributed to the C=O stretching of the carboxylic group, and the peak at 1600 cm^-1^ is attributed to the N-H group bending mode. A sharp peak at 1405 cm^-1^ was attributed to the bending of the C-H group. The XPS survey spectrum (**Figure 1d)** confirmed the presence of carbon (C), nitrogen (N), sulphur (S), and oxygen (O) atoms on CNDs surface. The high resolution deconvoluted C1s spectrum consists of three peaks with the binding energies at 284.6 eV, 286.1 eV, and 289.1eV **(Figure S2a)**. These are assigned with the C-C, C-O-C, and O-C=O bonds, respectively. Similarly, the N1s spectrum deconvoluted into three peaks at binding energies of 399.66 eV, 400.72 eV, and 401.63 eV **(Figure S2b)**. These are ascribed to the C=C-NH_2_, R_2_-NH, and O=C-N-C. The S2p spectrum is deconvoluted into two peaks with binding energies of 168.8 eV and 169.97 eV, corresponding to the C-S and S-S **(Figure S2c)**. The O1s spectrum deconvoluted into the two peaks corresponding to the C-O and C=O, respectively **(Figure S2d)**. The elemental analysis showed that CNDs contained C (50.04%), O (31.86%), N (9.87%) and S (8.22%). Both the FTIR and XPS data showed that S-H, NH_2_, and COOH functional groups are predominantly remaining on the CNDs surface.

**Figure 1:**
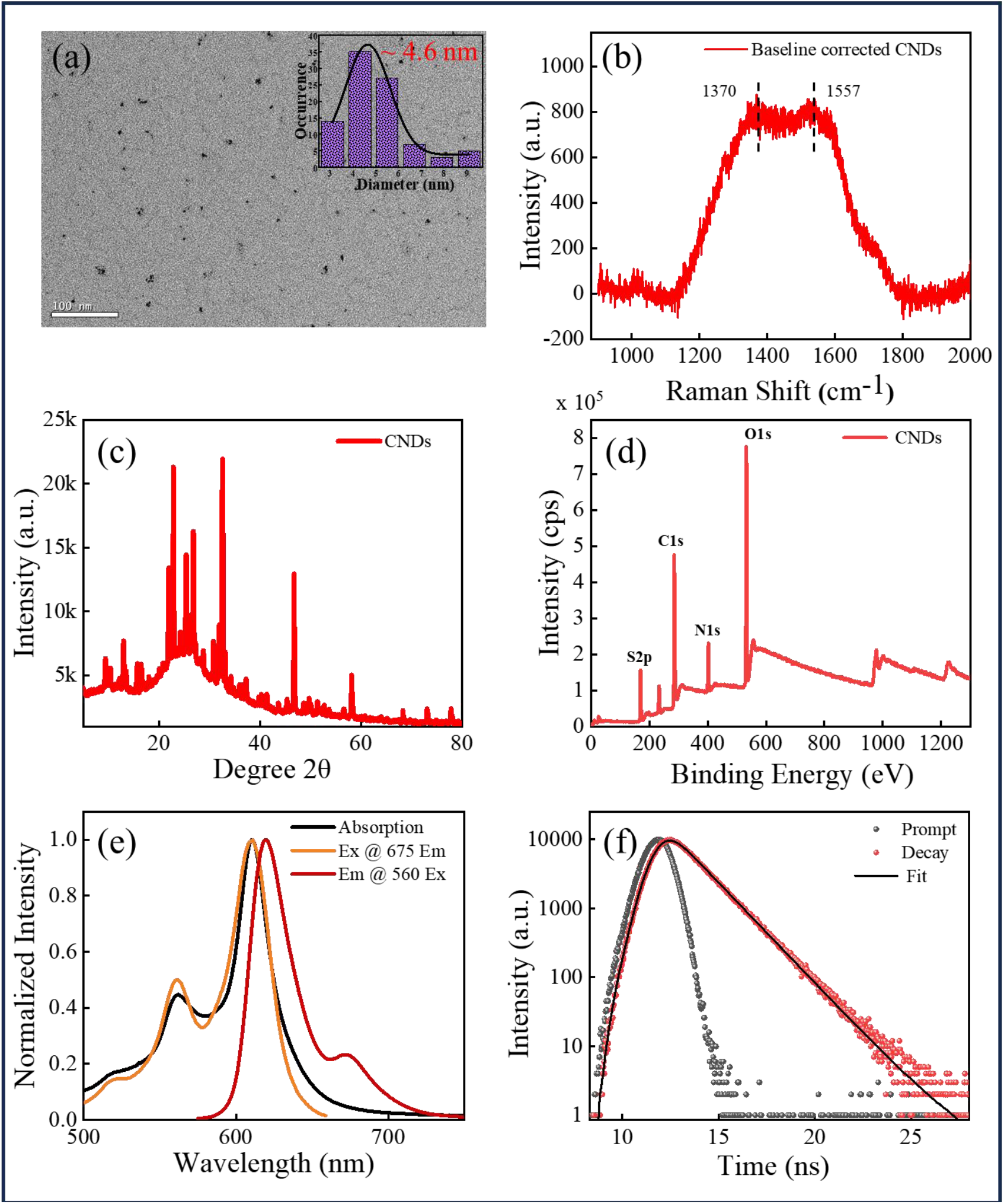
Characterization of CNDs: (a) TEM image of CND: Inset of a is showing the average particle size is ∼4.6 nm. (b) Raman spectra is showing the typical signature of the D and G bands. (c) Powder XRD pattern is showing the broad band at around 24° due to the amorphous nature of CNDs. Some crystalline nature is also visible on the top of the broad spectrum (d) XPS spectrum is showing the presence of C, O, N and S atoms (e) Absorbance (black), excitation (orange), and emission (red) spectra. A high overlap of absorption and excitation spectra was observed (f) The fluorescence lifetime decay of CNDs was collected by capturing emission at 625 nm with an excitation at 574 nm. The obtained average lifetime was 1.5 ns.

The measured absorption spectrum of the CNDs showed a well-defined progression of the vibrational features with a peak maximum at 610 nm and 1^st^ and 2^nd^ overtone bands at 517 nm and 560 nm. These peaks suggest that apart from the core and edge state absorptions, CNDs also have molecular states like absorption^[36]^. The emission spectrum of the CNDs showed intense peaks at 625 nm and a shoulder peak at 675 nm, respectively, upon excitation at 560 nm (**Figure 1e**), which suggesting the red emissive nature of CNDs. The measured excitation spectrum, upon keeping the emission maximum fixed at 675nm, exclusively overlapped with the absorption spectrum. These data confirms that the emission in the red region originates from the absorbed molecular state species present on the surface of the CNDs. The fluorescence lifetime of CNDs was obtained as 1.5 ns when excited at 574 nm pulsed nano LED source, and the emission was collected at 625 nm (**Figure 1f)**. It is to be pointed out that CNDs typically showed excitation wavelength dependent emission spectra. However, the measured excitation wavelength dependent emission spectra (**Figure S3**), showed an excitation wavelength independent emission spectral feature in the red region above 600 nm. A little excitation dependency in emission spectrum was obtained with two major peaks at around 400 and 460 nm. The shorter wavelength peak is assigned as the core state emission, while the peak at around 460 nm is assigned to the surface state emission.

The cytotoxicity was performed by XTT assay, which showed that over 80% of HeLa cells survived at a dosage of 15 μg/mL with 24 h incubation. This result suggested very good biocompatibility of CNDs for biological application **(Figure S4a)**. We also measured the reactive oxygen species (ROS) and found no discernible amount of ROS generation as compared to the control experiment **(Figure S4b)**. Next, we carried out a cell internalization experiment in HeLa cells before continuing with real bioimaging application of CNDs. **Figure 2a** shows the confocal images of cells incubated with CNDs for 2 h. It is clear from the figure that CNDs have the excellent capability of staining both the nucleus and the cytoplasm but at 561 nm and 639 nm excitation lasers. An appropriate dichroic mirror and bandpass filter were used to collect emission in the respective orange and red channels. Considering that the CNDs stain the DNA inside the nucleus, a colocalization experiment was performed with DAPI **(Figure 2b i-iii)**. Pearson correlation coefficient (PCC) was calculated to be ∼0.8, which is strongly indicating that the CNDs stain the DNA inside the nucleus. We further carried out the DNA and RNA digestion studies using deoxyribonuclease enzyme (DNase) and ribonuclease enzyme (RNase), respectively, to confirm the DNA specific labelling of CNDs but not RNA. The DNase-treated HeLa cells (**Figure 2c)** showed the complete loss of the red emission, while RNase treated cell (**Figure 2d)** showed no change in fluorescence intensity. Thus, the RNA and DNA digestion test confirms that the CNDs binds explicitly to DNA only in the nucleus. We further explored the potential staining of CNDs in plant cell nucleus. we took the root tip of the tomato plant and incubated with CNDs for 1 h and then performed the confocal image. **Figure S5** shows that the CNDs efficiently bind to the nucleus of the tomato plant roots. To understand the cytoplasmic organelle specificity, we carried out several colocalization experiment and finally observed that CNDs specifically bind to the mitochondria. The measured colocalization experiment with Mito Tracker Green (MTG) showed a PCC value of 0.82 thus suggesting that CNDs stain the mitochondria in the cytoplasm (**Figure 2e**).

**Figure 2:**
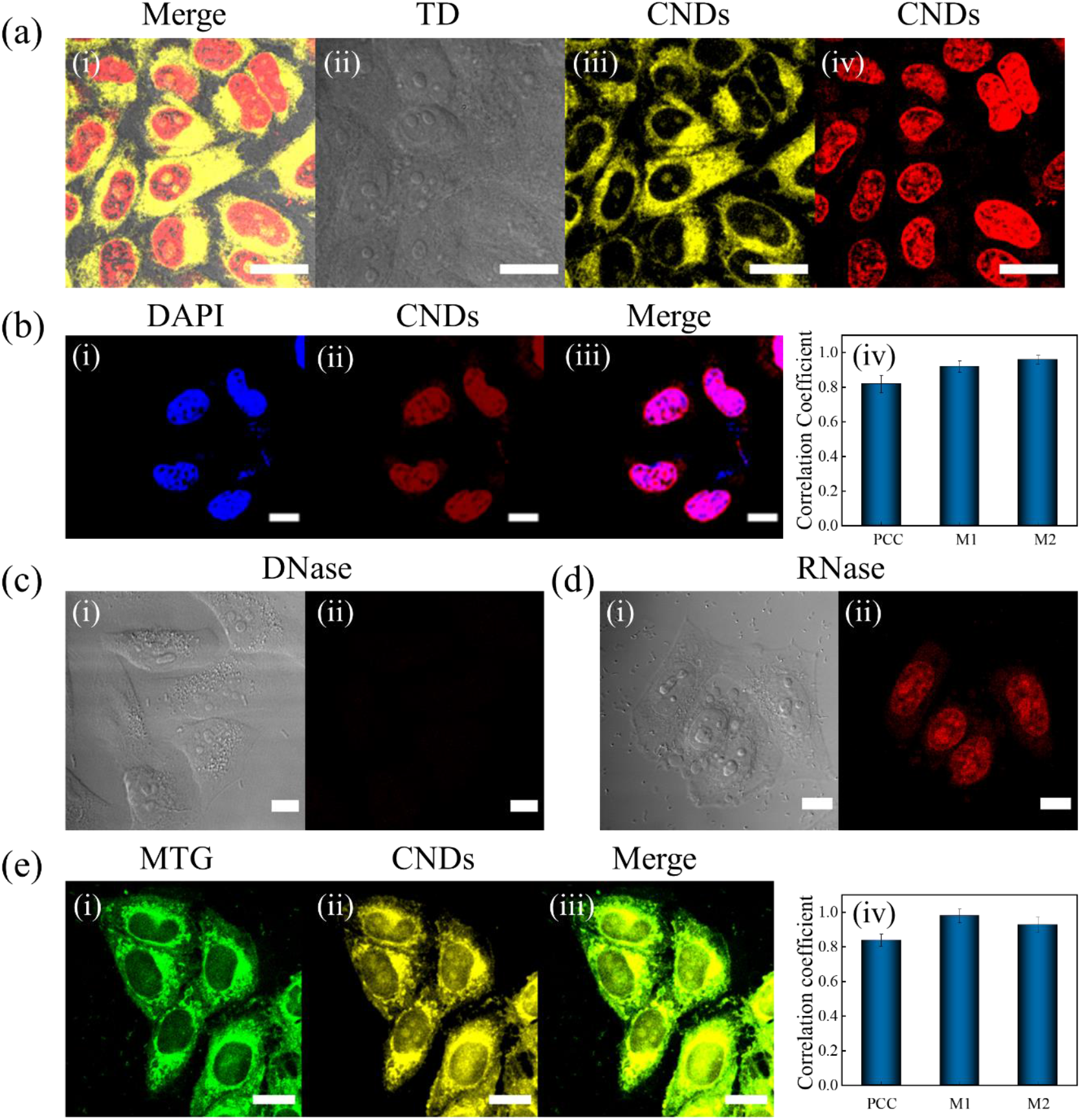
Cellular uptake and colocalization of CNDs: (a) Cellular internalization in HeLa cells suggests that CNDs stain both nucleus (red) and the cytoplasm (yellow). The red color emission was obtained when excited with 639 nm and emission was collected with 700/75 nm bandpass filter. The yellow color emission was obtained when excited with 561 nm laser, and the emission was collected with 595/50 nm bandpass filter laser (Scale bar: 20 µm). Here column i represent the merge image of transmission detection image (TD), and CNDs-stained nucleus and cytoplasm, ii represent TD image, iii & iv represent CNDs-stained cytoplasm and nucleus. (b) (i) DAPI (blue) staining and (ii) CNDs (red) staining along with (iii) the merged image shows the high colocalization of i and ii, (iv) shows the PCC and Mander’s overlap coefficients, M1 and M2. PCC, M1 and M2 coefficients with value >0.8 confirms the significant staining of CNDs to the nuclear DNA (scale bar: 10 µm). (c) DNase and (d) RNase digestion tests confirm that CNDs binds to the DNA within the nucleus but not RNA (scale bar: 10 µm). Columns i and ii represent TD and CNDs-stained nucleus. (e) Colocalization images of commercial MitoTracker Green (green color) with CNDs (yellow color), PCC value of 0.82 confirms the mitochondrial staining of CNDs in the cytoplasm (Scale bar: 20 µm). Here columns i and ii represent MTG and CNDs-stained mitochondria, iii represent their merged image (i & ii) and iv represent their corresponding correlation coefficients (PCC, M1 and M2).

Next, we observed the DNA dynamics in each step of the mitotic cell divisions using confocal and SRRF microscopy. Each of the PSFs created by the single molecules, in SRRF, contained a higher degree of local geometrical symmetry than the background. It calculates the local gradient convergence, termed as radiality, in the whole frame by dividing each pixel into sub pixels, thus preserving the information in the gradient field. The confocal and SRRF images of the nucleus using CNDs as a staining probe allow facile identification of mitotic cell phases in HeLa cells, as presented in **Figure 3** ^[37]^. Mitosis is a process of cell duplication in which parent cells give rise to two genetically identical daughter cells, each with the same number and kind of chromosomes as that of the parent cell. The complete process of mitosis consists of prophase, metaphase, anaphase, and telophase. Prophase is the initial stage of mitosis characterized by the condensation and compaction of chromatin into well-defined chromosomes, as shown in **Figure 3a**. In this phase, the nucleolus and nuclear membrane disappears. As prophase ends and metaphase begins, the microtubule pulls the chromosomes with equal force and aligns in the middle of the cell, known as the metaphase plate, as shown in **Figure 3b**. Before proceeding to anaphase, the cells check to ensure all the chromosomes are at the metaphase plate with their kinetochores correctly attached to microtubules. Metaphase leads to anaphase, during which each chromosome sister chromatid separates from each other and is pulled towards the opposite poles of the cell, as observed in **Figure 3c**. The centromere of each chromosome leads at the edge, where arms trail behind it. These sister chromatids become the chromosomes of each daughter’s nuclei. During the telophase, the chromosomes reach at the cell poles, and the mitotic spindle breaks down into its building blocks. Two new nuclei form, as shown in **Figure 3d**. The nuclear membrane and nucleoli also finally reappear at this stage, as is evident from the figure. The chromosomes begin to decondense and return to their stringy like form. Telophase is followed by cytokinesis, where the cytoplasm divides into two new daughter nuclei. The results presented in **Figure 3** strongly suggest that the CNDs are highly capable of staining the DNA in its coiled form and also in highly dense chromosomal form and are very efficient to observe each stage of the mitotic cell division.

**Figure 3:**
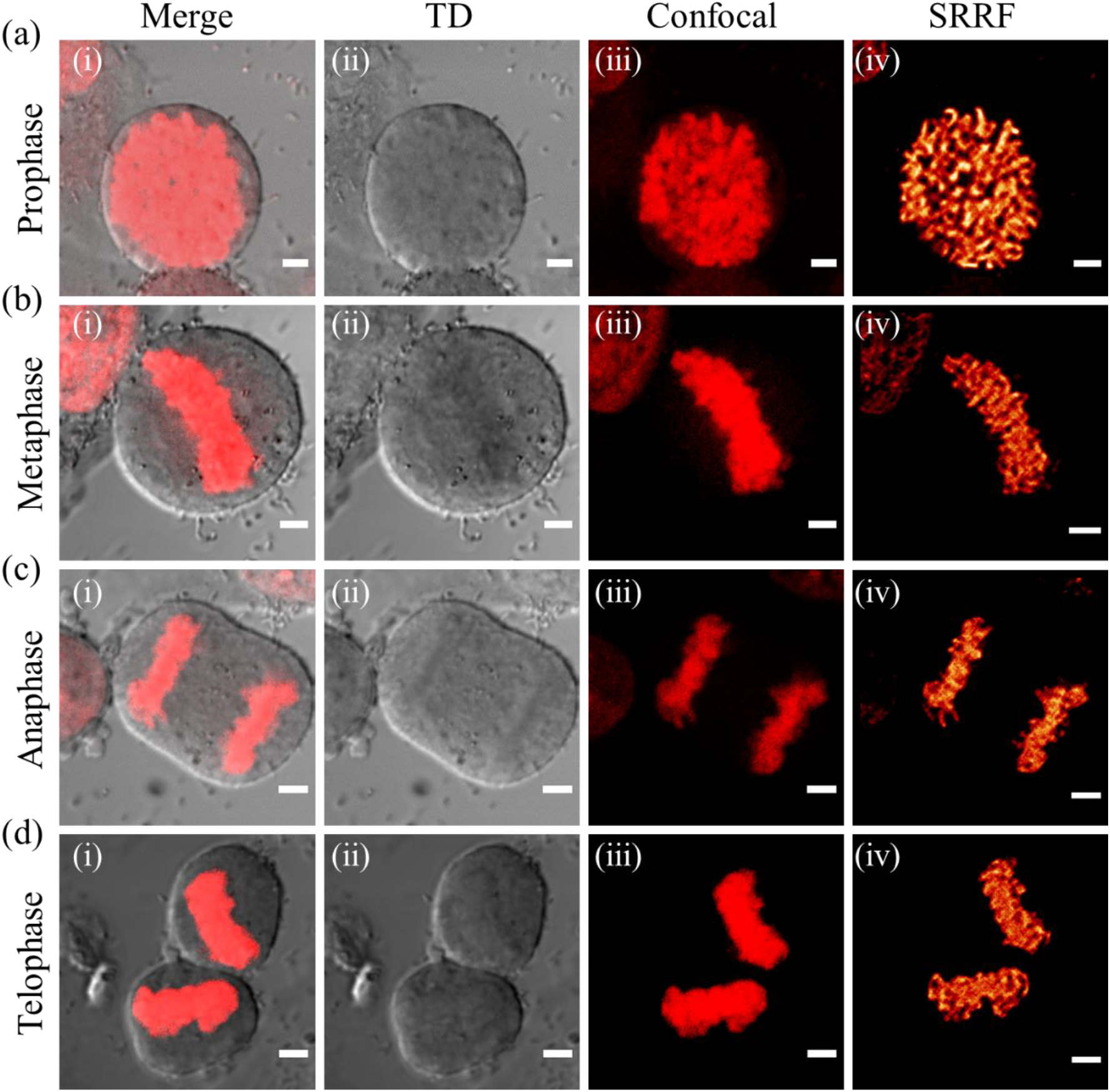
Confocal and SRRF Image of mitotic cell division in HeLa cells: Confocal and SRRF images of (a) prophase, (b) metaphase, (c) anaphase, and (d) telophase (scale bar: 3 µm). Columns (i), (ii), (iii), and (iv) represent the merge (TD & Confocal), TD, confocal and SRRF images respectively. A clear SRRF image of the chromosomes in each stage of the cell division was observed.

After completing all the necessary understanding of the DNA staining capability and the cell division using CNDs as a fluorescent probe, we next carried out the detailed experiment to understand the chromosome and chromatin dynamics upon paclitaxel treatment. Chromosomes in normal condition remained intact and condensed form while a significant fragmentation of the chromosomes was evident upon treatment of the HeLa cells with paclitaxel for 24 h (**Figure 4a & 4b**). We also observed the events of mis-segregated, lagging, and the bridging chromosomes in mitotic cell division at the anaphase state upon paclitaxel treatment (**Figure 4c-e**). This was not present in normal mitotic cell division. The statistical analysis (**Figure S6)** revealed that in normal conditions the average length of chromosomes is ∼2.5 µm while the average length of fragmented chromosomes after paclitaxel treatment is ∼1.5 µm.

**Figure 4:**
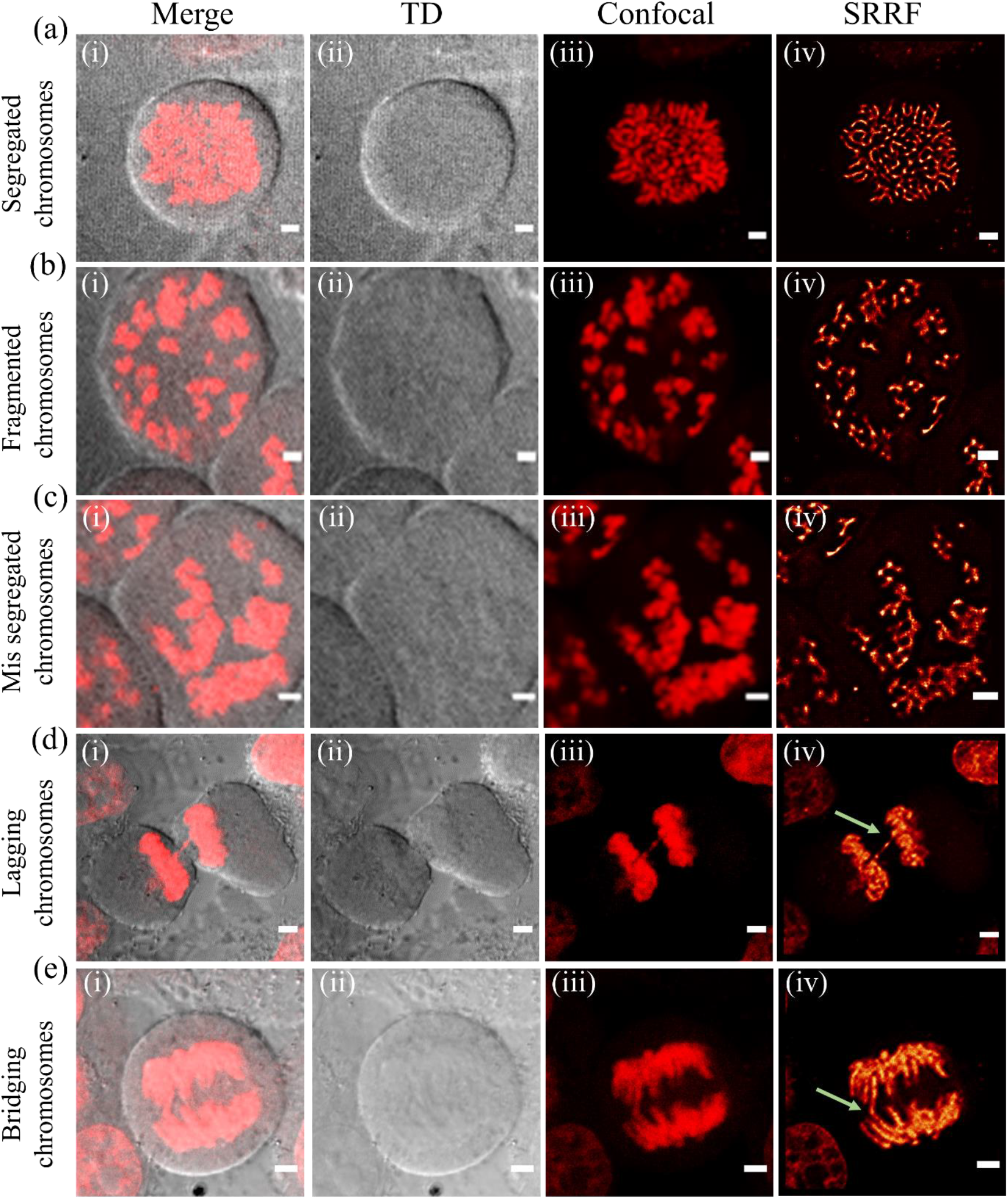
Nuclear chromosomal dynamics upon paclitaxel treatment. Confocal and SRRF images of CNDs stained in HeLa cells of (a) segregated chromosomes without paclitaxel treatment (b) fragmented chromosomes upon treatment with paclitaxel for 24 h and (c) mis-segregated chromosomes (d) lagging chromosomes (e) bridging chromosomes at anaphase stage upon treatment with paclitaxel for 3 h to 6 h. Columns (i), (ii), (iii), and (iv) represent the merge, TD, confocal and SRRF images respectively (scale bar: 3µm).

The above results inspired us to understand the detailed mechanism and pathway of action of paclitaxel. Hence, we carried out a time dependent paclitaxel treatment for 3, 6, 12 and 24 h and captured the DNA dynamics in each event. The SRRF image in **Figure 5a** showed that in normal conditions (without paclitaxel treatment), the DNA are widespread with the threat like structure throughout the nucleus and the outer nuclear membrane is also clearly visible (**Figure S7**). Interestingly, upon paclitaxel treatment within 3 h, the nucleus starts deforming in several instances where the membrane destabilizes, breaks down and finally disappears (**Figure 5b & Figure S8**). As a result, the normal segregation of chromosomes during cell division was disrupted and the DNAs were condensed in several segregated regions of the nucleus. As the nuclear membrane disappeared, the oversaturated SRRF image showed that the chromosomes are opened and formed few circular structures, which then again organised in several circular structure. (**Figure 5c & Figure S9**). A small nuclear membrane also formed in each case as the time progresses from 3 to 6 h. This result clearly suggested the formation muti-micronucleus (**Figure 5d & Figure S10**), which is followed by the lagging and mis-segregated chromosomes. These multi-micronuclei have several small nuclei rather than a single contiguous envelope which are clustered together. At 12 h, a complete muti-micronucleus was observed, which then fragmented at 24 h to process for the cell death (**Figure 5e & Figure S11**). These findings strengthen the anticancer activity through a stepwise mechanism for cytotoxicity of paclitaxel towards HeLa cells including breaking of the nuclear membrane, followed by the formation of lagging chromosomes, and finally, multi-micronucleus formation that leads to cell death. Hence, the current finding thus unambiguously demonstrates that CNDs is a potential fluorescent probe that captures all stages of the paclitaxel mechanism in anticancer activity with a strong support to non-mitotic pathway cell death during paclitaxel treatment.

**Figure 5:**
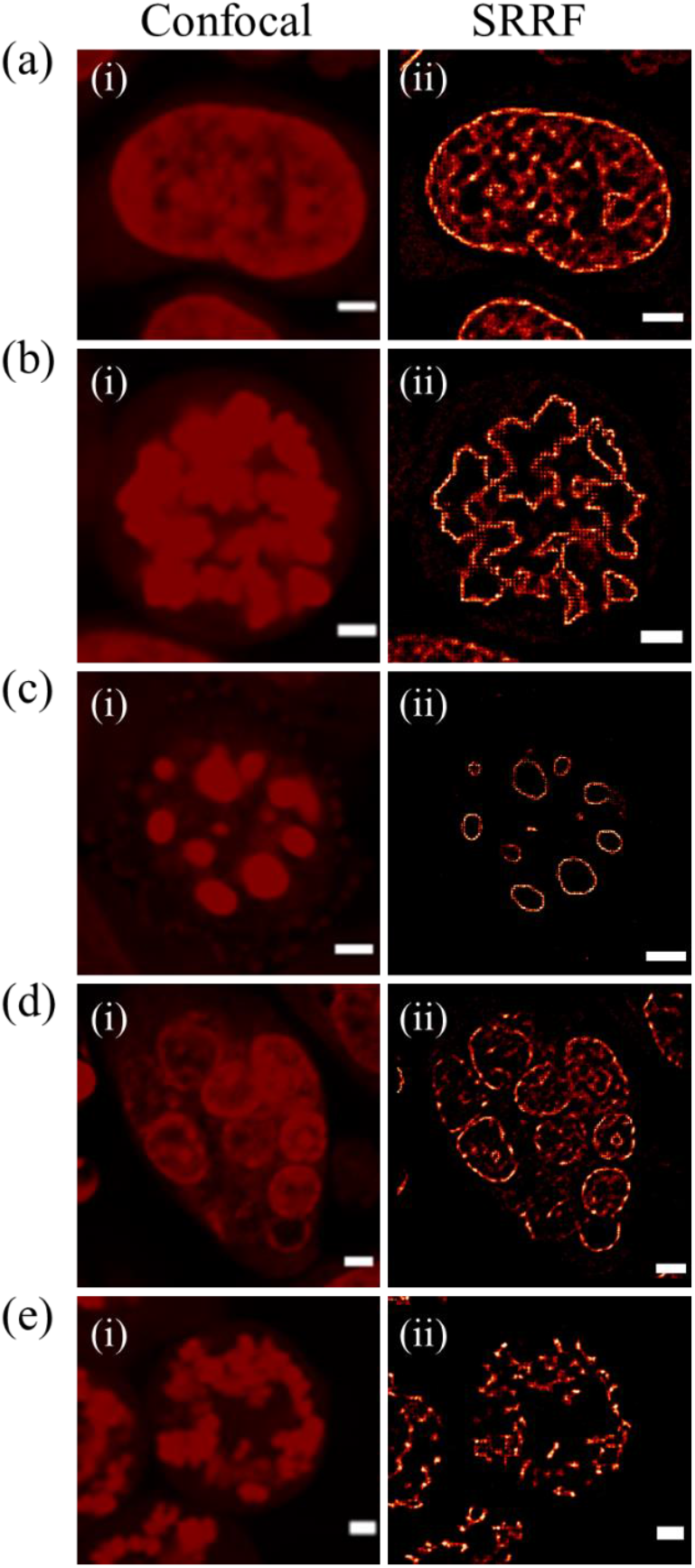
Time dependent DNA dynamics upon paclitaxel treatment. (a) The Nuclear DNA stained with CNDs in HeLa cell in the absence of paclitaxel. (b) after treatment with paclitaxel for 3 h disappearance of nuclear envelope followed by the chromatin chain opening and rearrangement. (c) the starting point of the formation of multi-micronucleus within 3 h to 6 h and continued till 12 h, that leads to d) the organised circular structure fully matured multi-micronucleus with dense DNA structure at 12 h. (e) spreading of chromosomes after breaking of the multinuclear membrane within 24 h. Columns (i), (ii), represent the confocal and SRRF images respectively (scale bar: 3 µm).

Finally, we explored the capabilities of CNDs to observe the chromatin dynamics as chromatin has been found to play a major role in nuclear and cellular function. Chromatin is tightly coiled DNA and histone proteins that constitute most of the nucleosome. A nucleosome is 147 bp of DNA wrapped in 1.67 left-handed twists around an H2A, H2B, H3, and H4 histone octamer. There are two types of chromatins: (a) heterochromatin and (b) euchromatin. Heterochromatin is a compact structure that is transcriptionally inactive and found in pericentric regions. On the other hand, euchromatin is less condensed gene-rich genomic regions and is more transcriptionally active. **Figure 6** shows the colocalization experimental data of CNDs with FITC-tagged commercial H3K4me3 and H3K9me3, which are very well known to bind with euchromatin and heterochromatin respectively. From the data and the calculated PCC value and confirmed that the CNDs colocalized with H3K9me3 efficiently but not with H3K4me3. These results suggested that CNDs specifically bind with heterochromatin but not with euchromatin. The specificity of CNDs for heterochromatin could be understood by their ability to bind to AT-rich regions of DNA, which are abundant in heterochromatin^[38,39]^. Getting inspired by the above results; we are keenly interested to understand the chromatin dynamics upon paclitaxel treatment. Therefore, a colocalization experiments of CNDs with H3K4me3 and H3K9me3 at multi-micronucleus stage and during the cell death processes were carried out. The PCC values for heterochromatin are quite significant (0.7) for heterochromatin shown in **Figure 7 & figure 8**, but for euchromatin it is only 0.4. This observation suggests the potential alternations in heterochromatin structure during the multi-micronuclei stage of paclitaxel treatment. The widespread distribution of euchromatin indicates their inactive participation in all successive stages during paclitaxel treatment. A report on wheat and pearl millet hybrid embryos showed that multi-micronuclei contained exclusively heterochromatin^[40]^. The study further proposed that the heterochromatinization and DNA fragmentation in multi-micronuclei formed by extrusion of paternal chromatin from interphase nuclei is involved in the pathway of haploidization. Our results of the chromatin dynamics during the multi-micronuclei stage of paclitaxel treatment suggested that heterochromatin played a significant role in the condensed stage of multi-micronuclei.

**Figure 6:**
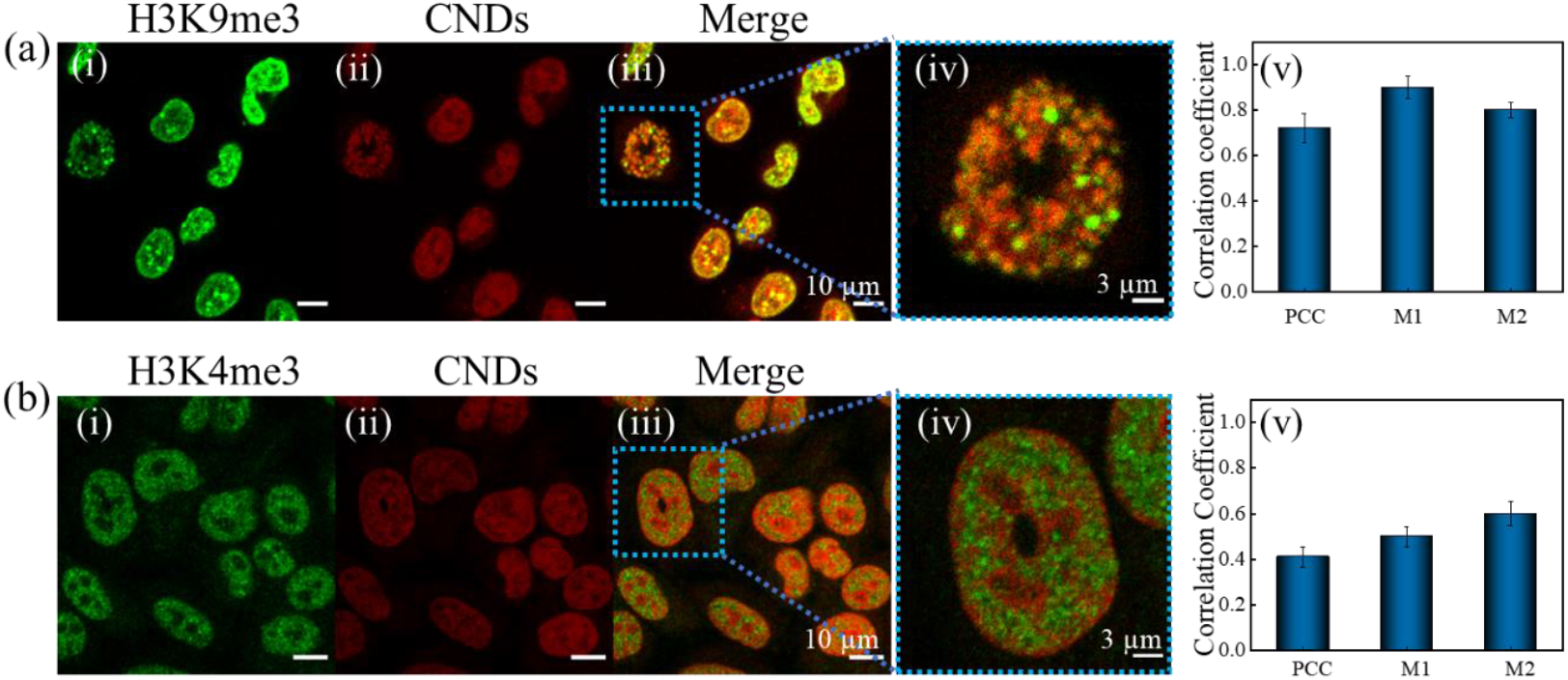
Colocalization of CNDs with H3K9me3 and H3K4me3: (a) Representative colocalization images of heterochromatin stained with H3K9me3 (Green) and CNDs (red). The PCC value was ∼0.74 suggesting significant colocalization in between H3K9me3 and CNDs in the nucleus. (b) Representative colocalization images of euchromatin stained with H3K4me3 (green) and CNDs (red). The PCC value was ∼0.40 indicating almost no colocalization between H3K4me3 and CNDs in the nucleus. Columns (i) represent staining with H3K9me3 (top) and H3K4me3 (bottom), (ii) represent CNDs staining, (iii) represent the merge of (i) and (ii), and (iv) represent the zoomed image of the marked area of (iii). (v) shows the respective correlation coefficients PCC, M1, and M2.

**Figure 7:**
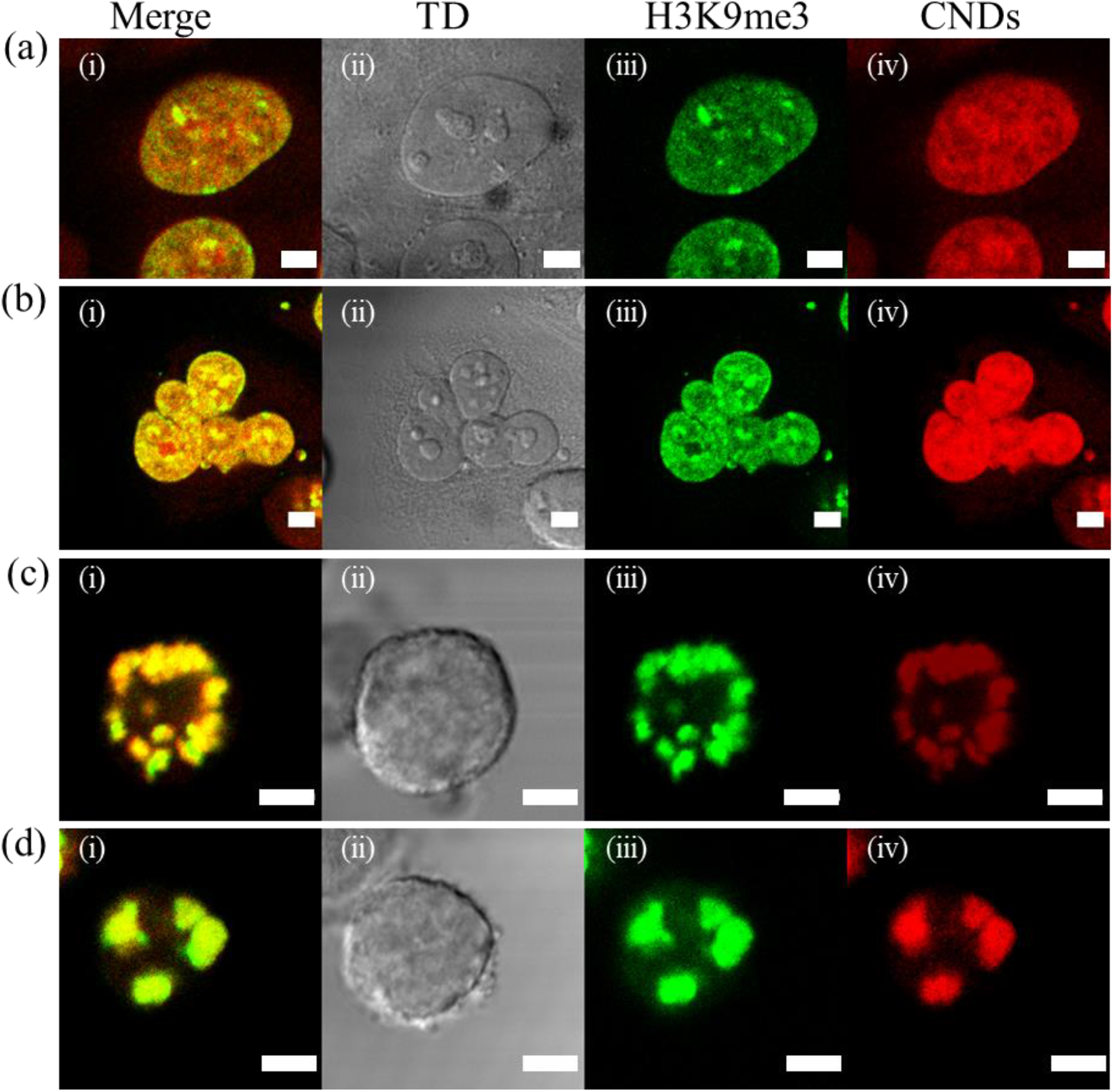
Colocalization of CNDs with H3K9me3 during paclitaxel treatment at different Successive stages: CNDs-labelled HeLa cell nuclei are shown in red. The HeLa cells were counterstained with H3K4me3 shown in green color. The yellow colour represents the merged image of H3K4me3 and CNDs-stained nucleus images. A significant overlap was observed where (a) showing the colocalization in normal conditions, (b) showing colocalization during multi-micronucleus formation, after paclitaxel treatment (c) during fragmented chromosomal condition upon paclitaxel treatment (d) during cell death. Column i, ii, iii and iv showing the merge, TD image, H3K4me3 and CNDs staining in HeLa cells respectively (Scale bar: 5 µm).

**Figure 8:**
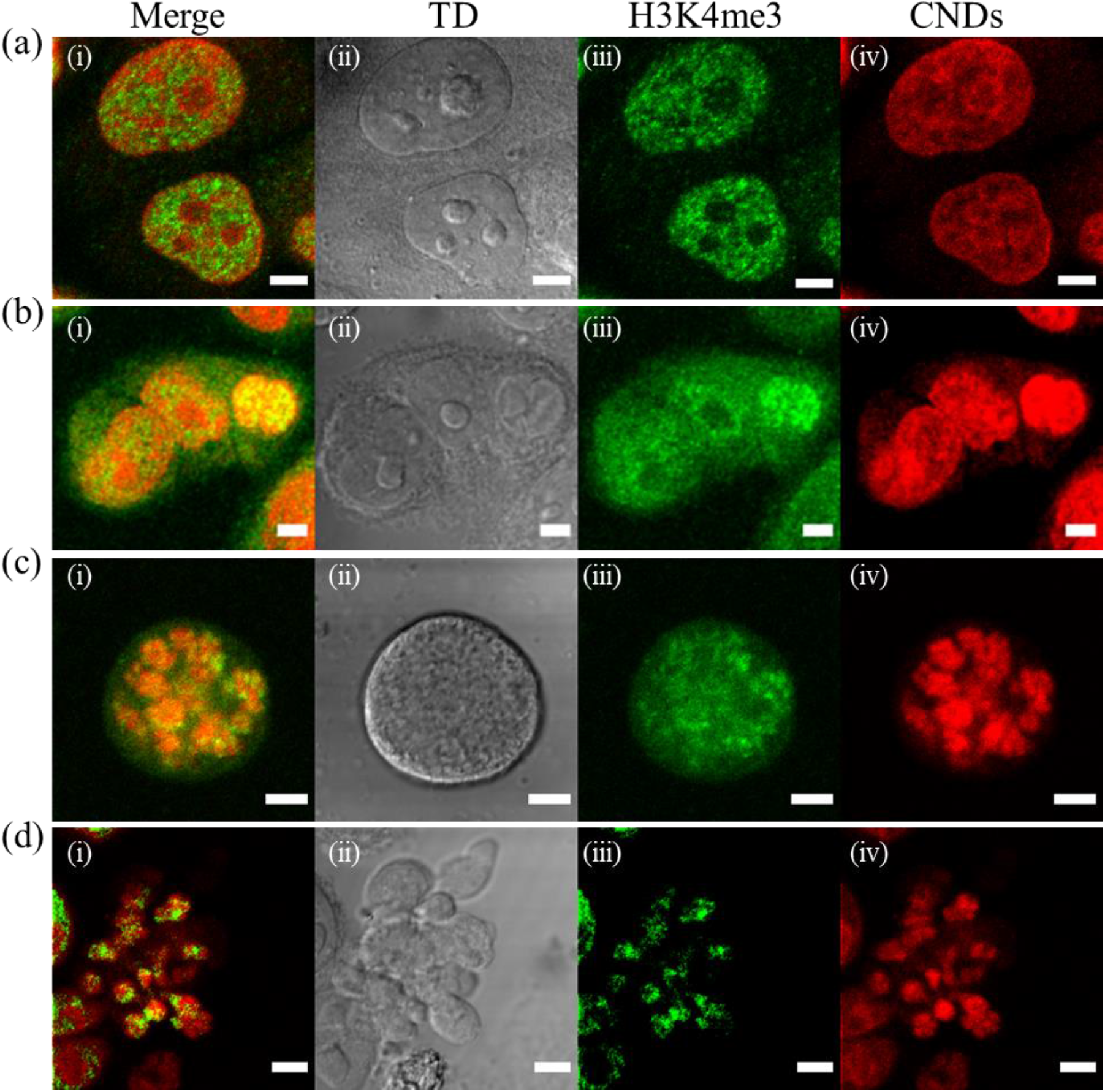
Colocalization of CNDs with H3K4me3 during paclitaxel treatment at different successive stages: CNDs-labelled HeLa cell nuclei are shown in red. The HeLa cells were counterstained with H3K4me3 shown in green color. The yellow colour represents the merged image of H3K4me3 and CNDs-stained nucleus images. Here a poor overlap was observed where (a) showing colocalization in normal conditions, (b) showing colocalization during multi-micronucleus formation, after paclitaxel treatment (c) during fragmented chromosomal condition (d) during cell death. Column i, ii, iii and iv showing the merge, TD, H3K4me3 and CNDs staining in HeLa cells respectively (Scale bar: 5 µm).

## Conclusion

In summary, using SRM along with synthesized non-toxic, biocompatible and highly fluorescent CNDs, we were able to visualize the nuclear DNA dynamics directly and unveiled the mechanism of paclitaxel treatment. CNDs successfully visualized the formation of lagging, mis-segregated, and bridging chromosomes that lead to multi-micronucleus formation. A detailed chromatin remodelling analysis suggested that heterochromatin played an important role during paclitaxel treatment. This insight into heterochromatin reorganization mechanisms will enhance our understanding of drug actions, offering a potential strategy to overcome chemotherapeutic resistance and improve the rational design of drug delivery for improved therapeutic efficacy of paclitaxel.

## Supporting information

Electronic Supplementary Information

## Data availability

All experimental and characterization data that support this article are available in the main manuscript and its ESI.

## Author contributions

RG designed and performed all bulk-level experiments with the help of RK. RG performed all cellular experiments. KK analysed the SRRF data with the help of AS and RG. SC performed plants staining experiment. RG wrote the article with the help of all co-authors. CKN supervised and guided the whole project.

## Acknowledgments

CKN thanks to the Science & Engineering Research Board (Project No. IITM/SERB/CKN/310) for the financial support. CKN is also acknowledges funding form the Council of Scientific & Industrial Research (IITM/CSIR/CKN/449). RG thanks the Prime Minister Research fellowship for the scholarship. KK and AS thanks the Ministry of Education, India, for their scholarships. CKN is thankful to the facilities provided by the Advanced Material Research Centre (AMRC), IIT Mandi. CKN thanks IIT Mandi for providing a state-of-the-art instrumentation facility including the cell culture.

