## Supplementary material for "Carbon nanodots as a red emissive fluorescent probe for the super-resolution microscopy of DNA dynamics during paclitaxel treatment": Electronic Supplementary Information

### for

### Materials and Methods

#### Chemicals and Materials:

O-phenylenediamine (OPDA) and Glutathione (GSH) were purchased from Sigma Aldrich and Thermo Fisher Scientific simultaneously. HCl was purchased from rankem. Dulbecco's Modified Eagle Medium (DMEM), 1% Antibiotic-Antimicotic, Penstrap, and fetal bovine serum (FBS) were purchased from Gibco. Paclitaxel was collected from Merk. Primary antibodies H3K4Me3 and H3K9Me3 were acquired from ABclonal. Secondary antibody (FITC conjugated secondary antibody) was also obtained from ABclonal. All chemicals were of analytical quality and did not require further purifications. All glassware and hydrothermal were washed with aqua regia (3HCl:1$HNO_3$), ethanol, acetone, and rinsed several times with double distilled water. Double-distilled (18.3 MΩ) deionized water (Elga Purelab Ultra, Vivendi water system Ltd, India) was used throughout the experimental process. The cell lines (HeLa) used in this study were purchased from the National Centre for Cell Science (NCCS) Complex, Pune, Maharashtra, India - 411 007.

#### Characterization Techniques:

#### UV-Visible spectroscopy

The UV-Vis of CNDs was measured on a (50 W halogen lamp (2000 h life)) Shimadzu UV-Vis 2450 spectrophotometer. For absorption spectra, samples were placed in a transparent quartz cuvette with 1 ml volume and 10 mm path length.

#### Steady-state fluorescence

Steady-state fluorescence was recorded using a Horiba Fluorolog-3 spectrofluorometer from 280 to 700 nm excitation wavelength range. Again, transparent quartz cuvettes with 1 ml volume were used to capture emission spectra of CNDs.

**Fluorescence lifetime**

The fluorescence lifetime was examined by using the Horiba scientific Delta Flex TCSPC system with 574 nm Pulsed LED Sources. Ludox has been used as an IRF. By bi-exponentially fitting, the photon decays in different channels with a chi-squared value < 1.1, the fluorescence lifetime was identified.

**Thermogravimetric Analysis (TGA)**

Thermal properties were measured by Perkin Elmer Pyris Thermogravimetric analyzer from 25 °C to 800 °C with a heating rate of 10 °C $min^{-1}$ under a nitrogen atmosphere.

**Transmission electron microscopy (TEM)**

Transmission electron microscopy measurements were obtained through FEI TECHNAI, USA, FP 5022/22-Tecnai G2 20 S-TWIN transmission electron microscope, which operates at 200 keV using $LaB_6$ filament as an excitation source.

**Fourier transform infrared (FTIR) spectra**

Fourier transform infrared (FTIR) spectra of dried CNDs, GSH, and OPDA were acquired by using a Perkin-Elmer FTIR spectrophotometer equipped with a horizontal attenuated total reflectance (ATR) accessory containing a zinc selenide crystal and operating at 4 $cm^{-1}$ resolution. Spectrum was recorded using Resolutions Pro FTIR software by subtracting background spectra from the sample. The wavelength range to record FTIR spectra of CNDs is 400-4000 $cm^{-1}$.

**Raman spectrum**

The Raman spectrum of dried CNDs was measured by the confocal microscope Raman spectrometer (Horiba Scientific, Xplo RA ONE) with a 532 nm laser (spectral range 400 $cm^{-1}$ to 3500 $cm^{-1}$).

**X-Ray Photoelectron Spectroscopy (XPS)**

Surface chemistry of the material and elemental composition is measured by using XPS in which Auger Electron Spectroscopy (AES) Module PHI 5000Versa Prob II, FEI Inc. and $C_{60}$ sputter gun have been used for the characterization and scanning the spectra for $C_{1S}$, $N_{1S}$, $S_{2p,}$ $O_{1S}$ region.

**Confocal Imaging of HeLa Cells**

**Coverslip preparation:** The glass slides and coverslips were cleaned by incubating in freshly prepared Piranha solution for 30 min and finally washing with 70% ethanol in bath sonication, then dried at room temperature.

**Cell Culture, fixation, and staining:** All the cell culture experiments and slide preparations were performed in compliance with the relevant guidelines and norms of biosafety level-1 requirements of the Indian Institute of Technology Mandi. Cell lines were maintained by following the recommended protocols of the National Centre for Cell Science (NCCS) Complex, Pune, Maharashtra, India - 411 007.

Human cervical cancer (HeLa) cell lines were cultured in Dulbecco's Modified Eagle Medium (DMEM) with 10% fetal bovine serum (FBS), 1% anti-anti, and penicillin/streptomycin at 37 °C with 5% $CO_2$ humidity. The cells were grown in a 6-well plate on coverslips with $10^4$ cells per 100 µl density. Each well was filled with 2 ml of growth medium, and the cells were allowed to grow overnight for proper adherence and growth. Cell growth and attachment to the coverslips were examined with an optical microscope. Once the cells reached the proper adherence and confluency, they were stained with synthesized CNDs fluorescent probe to achieve enough labeling density for confocal microscopy. Finally, the cells were fixed by incubating with 4% paraformaldehyde solution in 1X PBS buffer for 15 min. The fixed cells were then washed 2-3 times with PBS buffer to remove extra CNDs and Cell culture medium. The coverslips were fixed with the help of mounting media (9 glycerol: 1 1X PBS) on a glass slide before imaging.

For paclitaxel mechanism elucidation, HeLa cells were again treated with 1 µM paclitaxel for 3, 6, 12, and 24 h. After that, cells were washed with PBS, and again, fresh media was added and incubated with CNDs followed by washing and fixing in 4% paraformaldehyde. The fixed cells were washed 2-3 times with 1X PBS buffer to remove extra CNDs and Cell culture medium. The coverslips were fixed with the help of mounting media (9 glycerol: 1 1X PBS) on a glass slide before imaging.

**ROS detection:** ROS detection was achieved by seeding the cells (40 X $10^3$ cells per well). After the proper adherence of cells, they were incubated with CNDs for 24 h. After the incubation, 1µM of $H_2$DCFDA was added to the cell medium and incubated for 30 minutes. Finally, the fluorescence intensity of cells was measured at 520 nm after excitation with 490 nm, and identification of ROS was achieved and compared with the control sample.

**Cell cytotoxicity:** Cell cytotoxicity was detected with the help of an XTT kit (Roche XTT kit II). HeLa cells were seeded in 96 well plates and cultured overnight with a density of 4 $\times 10^3$ cells per well. Then cells were incubated with CNDs for 2 h, at 37 °C in a $CO_2$ incubator. After the incubation with CNDs, the XTT labeling mixture was added and further incubated for 12-16 h. Then, the absorbance of the sample mixture was measured by a microplate reader at 550 nm of wavelength. The reference wavelength was set at 650 nm and then analyzed the cell viability against CNDs with comparison to blank and control samples. The mean and standard deviation were calculated from five wells tested in parallel.

**Immunofluorescence staining of HeLa cells:**

For Immunofluorescence imaging, HeLa cells were grown in their standard medium. Cells were harvested using 1X Trypsin-EDTA, and seeding on coverslips with density of $1\times10^5$ cells per well. After ensuring proper adherence, cells were proceeded for Immunofluorescence staining. Prior to antibody staining, cells were fixed and permeabilized with 4% paraformaldehyde and 0.1% Triton X-100 for 10 minutes followed by three washing with 1X PBS. Then cells were blocked using 1% BSA in 1X PBS buffer as a Blocking buffer. Subsequently, HeLa cells were incubated separately with primary antibody (1 H3K4Me3 :100 1X PBS buffer) specific for euchromatin, and a similar concentration of primary antibody was used for H3K9Me3 which is specific for heterochromatin staining. Both antibodies were incubated for the overnight at 4 °C. The next day, cells were again washed with 1X PBS. HeLa cells were further incubated with 0.1% Tween 20 for 10 minutes followed by incubation of secondary antibody (FITC conjugated secondary antibody) for 2 h. Finally, the cells were incubated with 15 µg/ml CNDs to check their proper colocalization. Afterwards, cell containing coverslips were affixed onto a glass slide using a drop of mounting media and used for confocal imaging.

**Confocal microscopy:** Confocal imaging of HeLa cells was performed using Nikon Eclipse Ti inverted microscope and images were acquired using NIS-Elements software. The cell samples were excited by using both 560 nm and 639 nm lasers. Finally, the images were collected by choosing a proper filter set.

**SRRF Imaging**

We acquired multi-frame confocal images consisting of 500 frames in a stack, utilizing a 60X oil immersion objective on a Nikon Ti inverted microscope operating in continuous mode during time lapse. Videos were captured using Nikon's NIS-Elements software using pinhole size:1 AU, scan size:128 x 128 pixels, scan speed:4 fps, line averaging: disabled, Channel Series: none. The videos were processed for SRRF using the NanoJ-Core and NanoJ-SRRF plugins of ImageJ, which are available at https://sites.imagej.net/NanoJ-SRRF/plugins/ . We developed an ImageJ macro language script to perform batch SRRF analysis on a high-performance GPU enabled computer (with NVIDIA GeForce RTX 3070). We used default settings with a ring radius of 0.5, a radiality magnification of 5, and axes in ring 6. All videos were first drift-corrected using the ImageJ NanoJ-Core plugin.

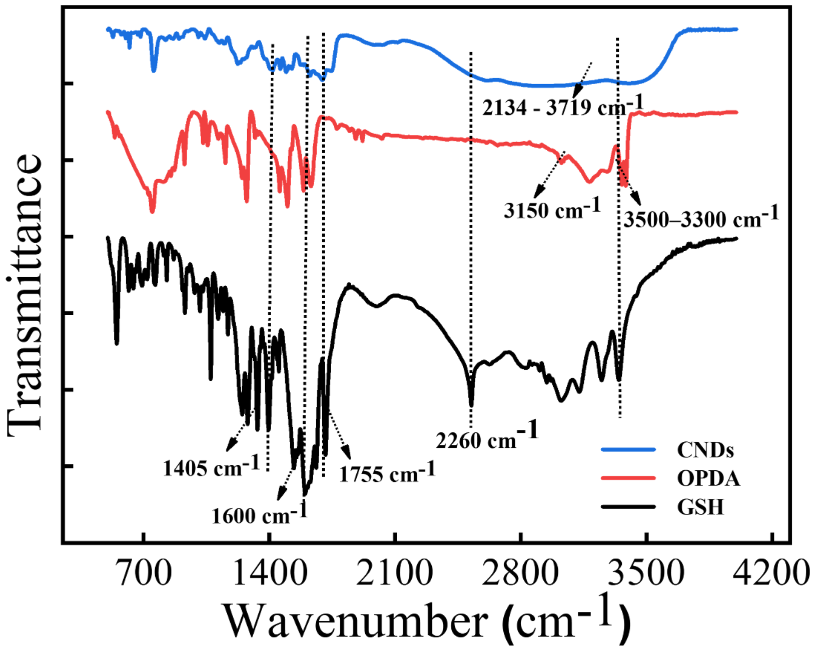


**Figure S1:** A comparison of the FTIR spectra of CNDs (Blue) along with precursor molecules OPDA (red) and GSH (Black).

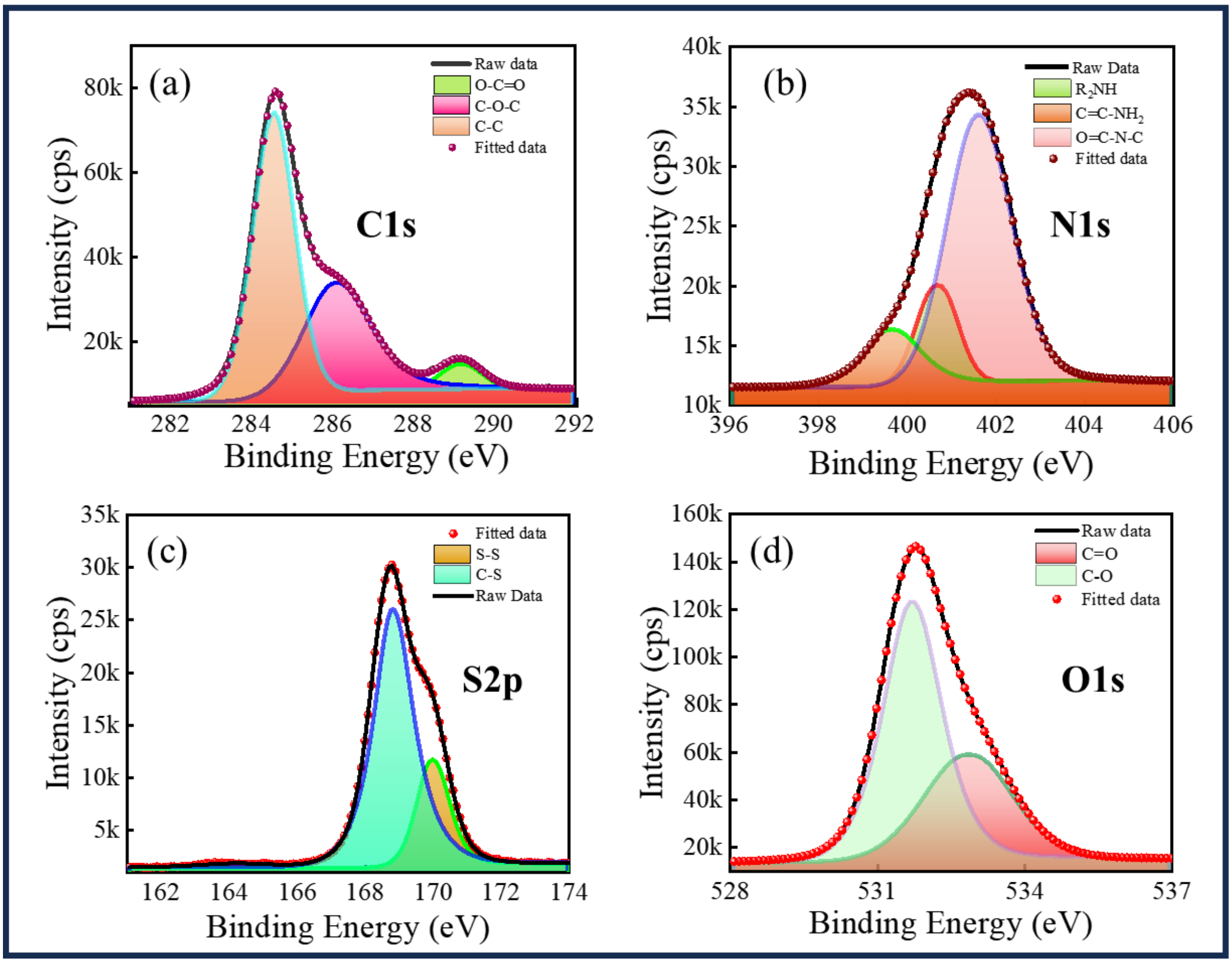


**Figure S2:** High-resolution X-ray photoelectron spectroscopy (XPS) spectra with deconvolution of (a) C1s (b) N1s (c) S2p and (d) O1s.

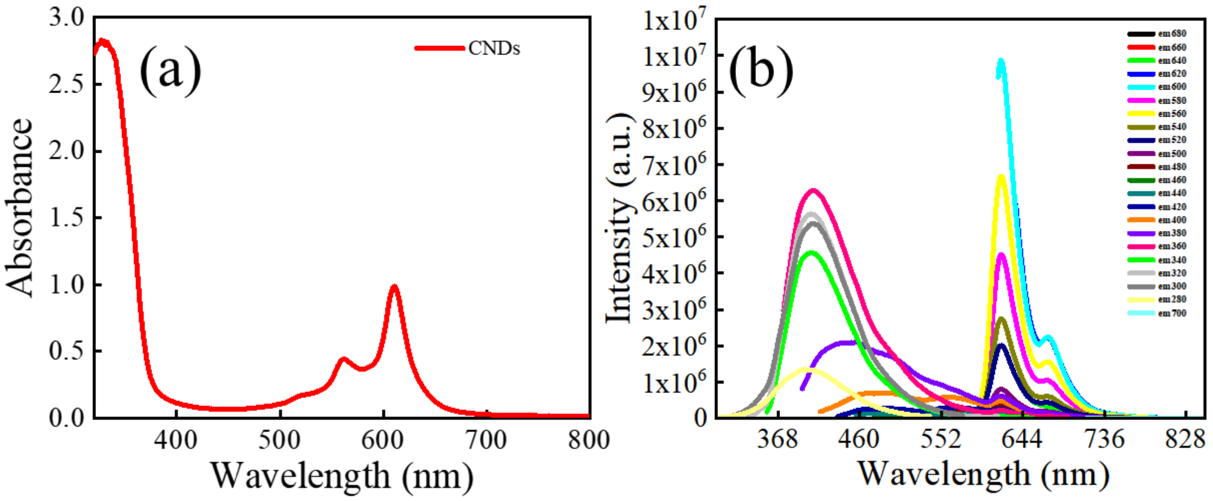


**Figure S3:** (a) Full UV-vis spectra of CNDs. (b) Excitation wavelength dependent fluorescence spectra of CNDs.

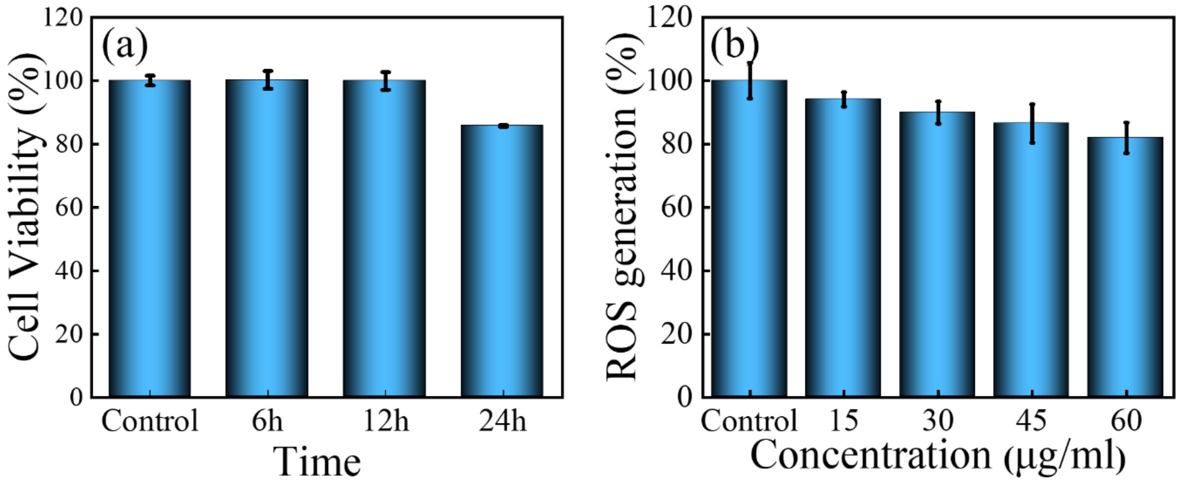


**Figure S4:** (a) Cell viability of HeLa cells incubated with 15 µg of CNDs at 37 °C for 6h, 12h, and 24 h. More than 80 % cell viability was observed after 24 h thus suggesting very less toxicity of the CNDs (b) Intracellular ROS generation in HeLa Cells showing very less ROS production

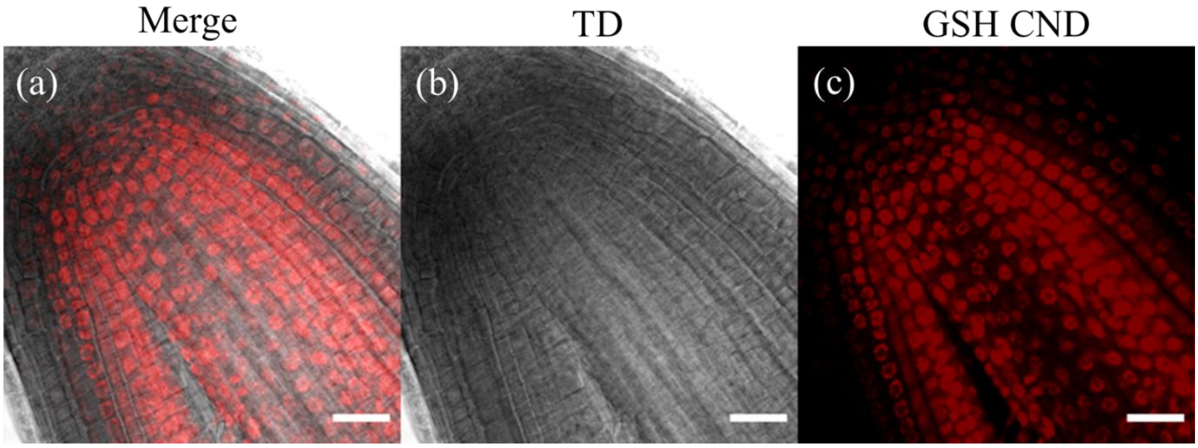


**Figure S5:** Confocal image of *Solanum lycopersicum* (tomato) roots tip stained with CNDs (a) display the merged image of b and c. (b) TD image (c) CNDs-stained nucleus. (scale bar: 30 μm)

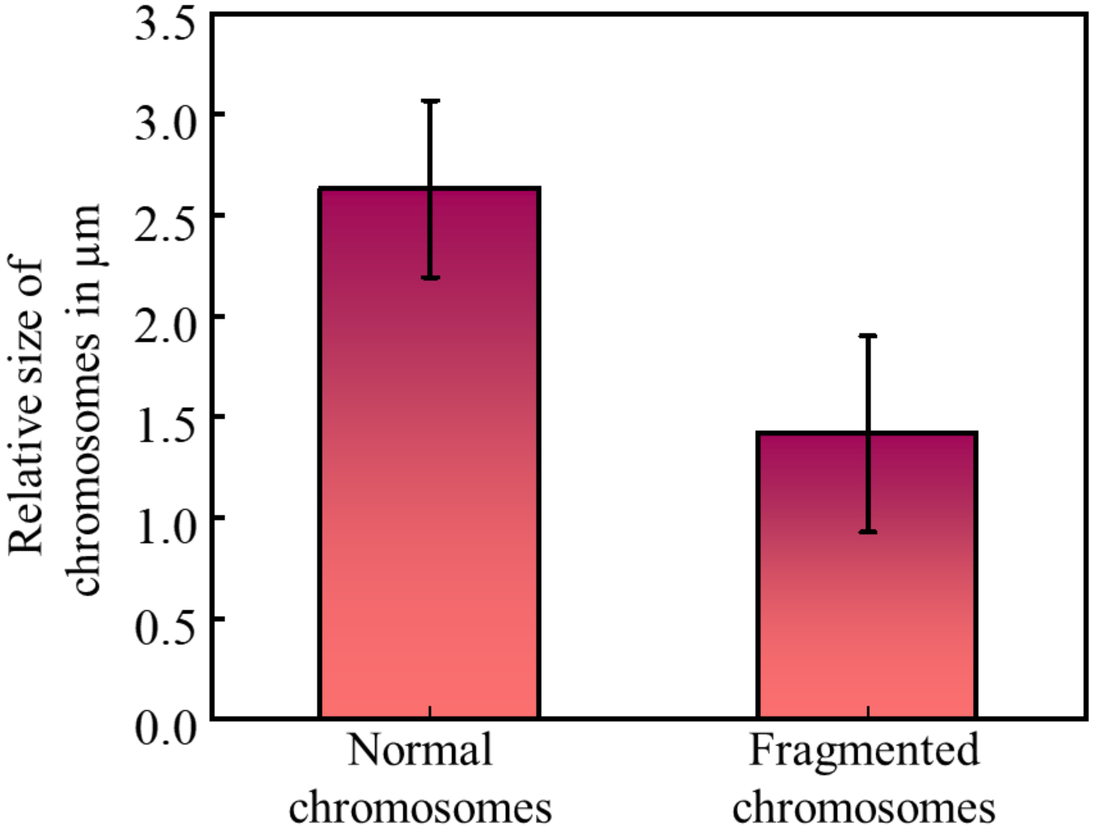


**Figure S6:** Statistical size analysis of chromosomes in normal conditions and after 24 h paclitaxel treatment.

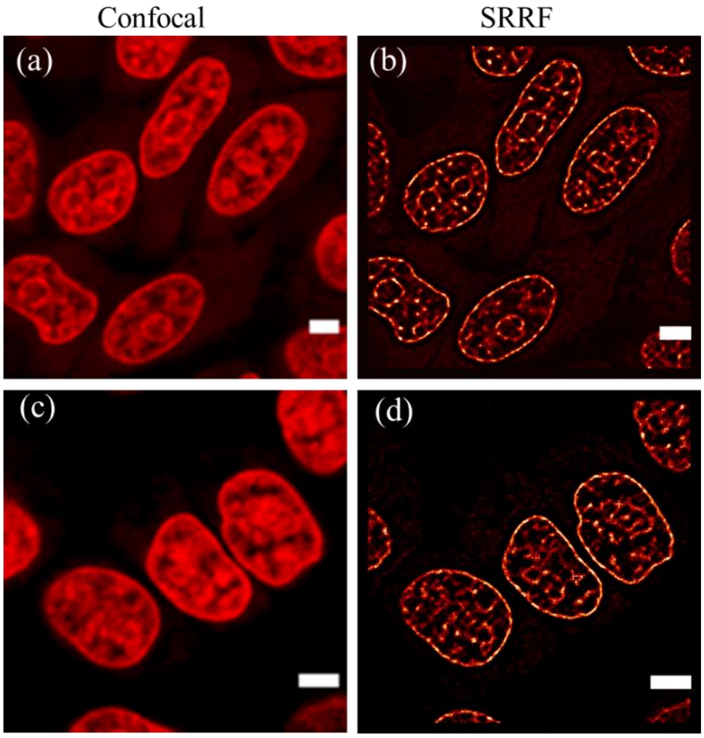


**Figure S7:** Confocal and SRRF images of HeLa cell nucleus stained with CNDs in the absence of paclitaxel (scale bar: 5 μm). In normal conditions, each cell is uninucleated showing single, intact nucleus surrounded by continuous nuclear envelope. Here a & c represent confocal images, b & d represent SRRF images.

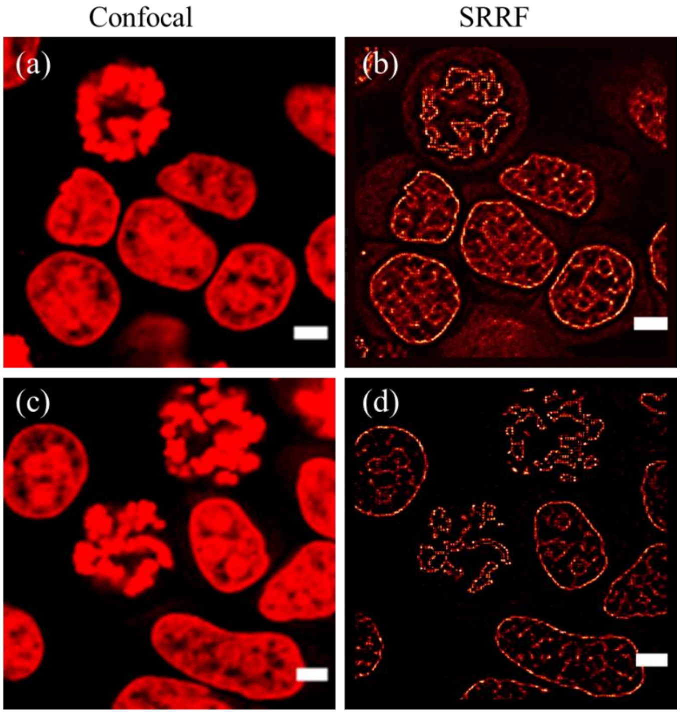


**Figure S8:** Confocal and SRRF images of HeLa cell nucleus stained with CNDs after 3 h treatment with paclitaxel (scale bar: 5 µm). The nucleus begins to deform and the nuclear membrane gradually start to disappear. Here a & c represent confocal images, b & d represent SRRF images.

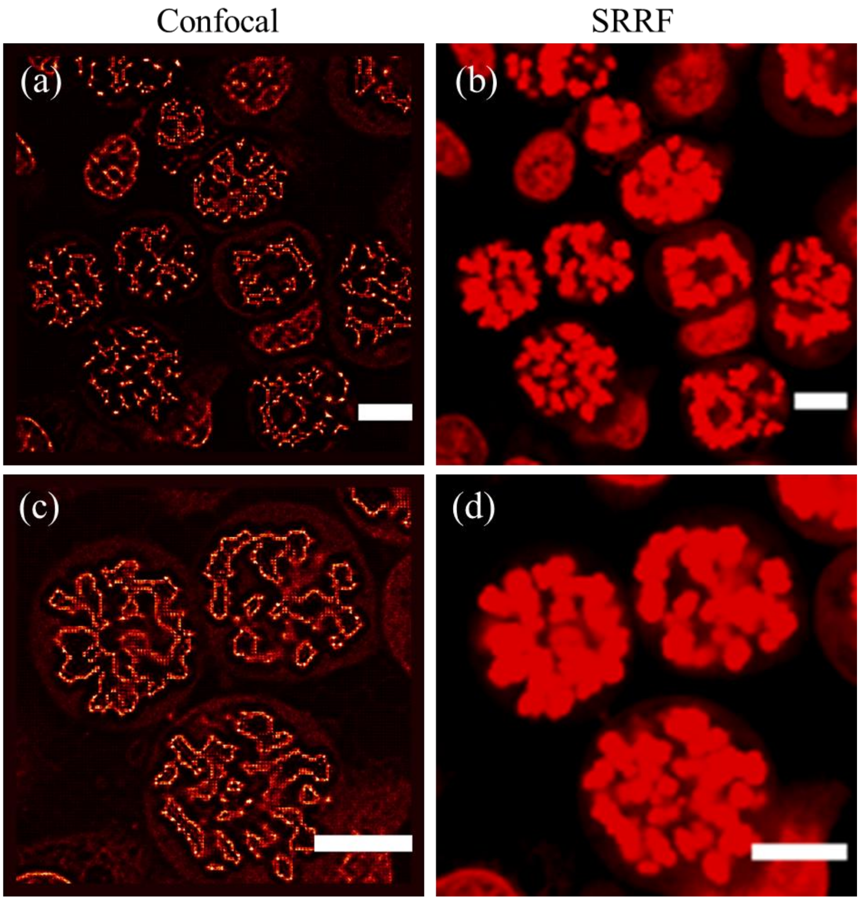


**Figure S9:** Confocal and SRRF images of HeLa cell nucleus stained with CNDs after 6 h treatment with paclitaxel (scale bar: 10 μm). After nuclear membrane disappearance, mostly cells undergo rearrangement of nuclear chromatin which lead to the formation of multimicronucleus. Here a & c represent confocal images, b & d represent SRRF images.

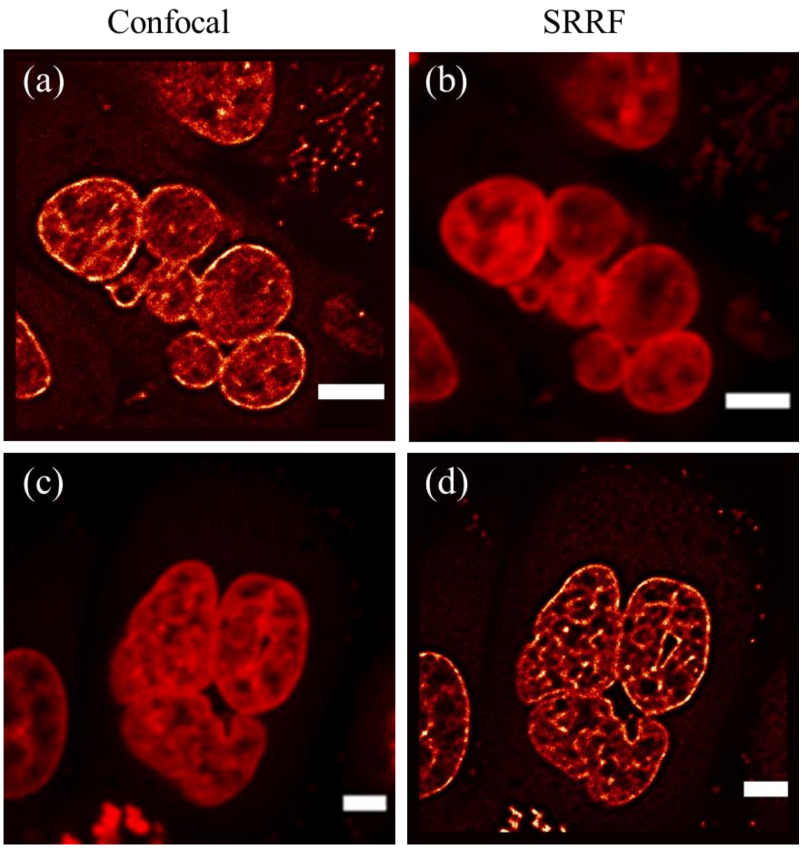


**Figure S10:** Confocal and SRRF images of HeLa cell nucleus stained with CNDs after 12 h treatment with paclitaxel (scale bar: 5 µm). Complete multi micronucleus formed and each nuclei form its own nuclear envelope. Here a & c represent confocal images, b & d represent SRRF images.

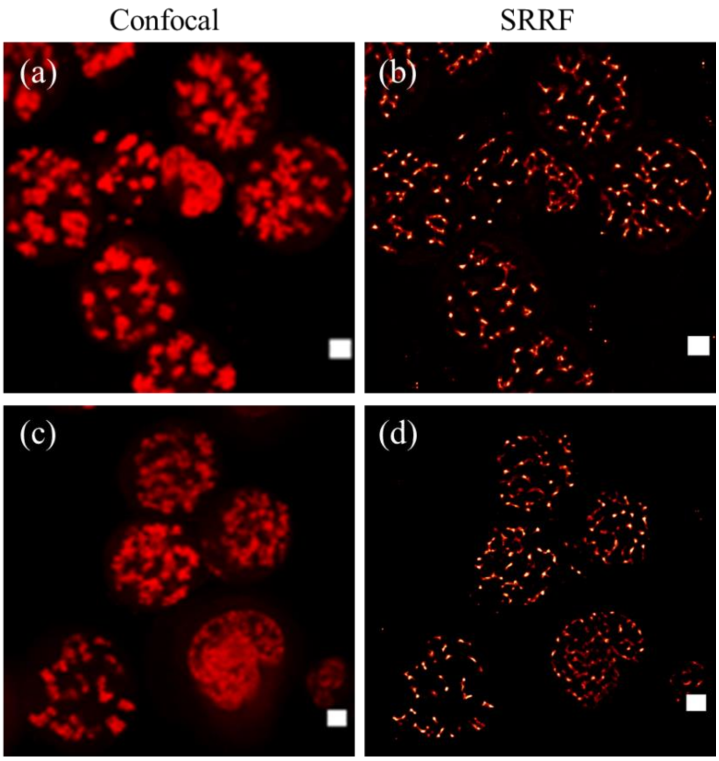


**Figure S11:** Confocal and SRRF images of HeLa cells nucleus stained with CNDs after 24 h treatments with paclitaxel. Multi-micronucleus containing fragmented and mis-segregated chromosomes are excluded into the cytosol (scale bar: 5 µm). Here a & c represent confocal images, b & d represent SRRF images.
